# Biological and experimental factors that define the effectiveness of microbial inoculation on plant traits: a meta-analysis

**DOI:** 10.1101/2024.04.30.591815

**Authors:** Hamed Azarbad, Robert R. Junker

**Affiliations:** Evolutionary Ecology of Plants, Department of Biology, University of Marburg, Karl-von-Frisch-Strasse 8, 35043, Marburg, Germany

**Keywords:** Bacteria, Inoculation, Experimental setting, Fungi, Microbes, Plant traits

## Abstract

Bacterial and fungal microbiomes associated with plants can significantly affect the host’s phenotype. Inoculating plants with one or multiple bacterial and fungal species can affect specific plant traits, which is exploited in attempts to increase plant performance and stress tolerance by microbiome engineering. Currently, we lack a comprehensive synthesis on the generality of these effects related to different biological (e.g., plant models, plant traits, and microbial taxa) and experimental factors. In a meta-analysis, we showed that the plant trait under consideration and the microbial taxa used to inoculate plants significantly influenced the strength of the effect size. In a methodological context, experiments under sterilized conditions and short-term periods resulted in larger positive effects on plant traits than those of unsterilized and long-term experiments. Based on our results, we propose a comprehensive checklist as a reference for future research to standardize the design, implementation, and reporting of microbial inoculation studies. We recommend that future studies should exploit the full range of the precision-realism continuum involving (short-term) lab experiments with sterilized plants and single inoculants but also and more often (long-term) field or greenhouse experiments with naturally occurring microbial communities associated with the plants and inoculated consortia including both bacteria and fungi.

## Introduction

Plants are colonized by a wide range of microbes that may have pathogenic, beneficial, or neutral effects on the host plant [1, 2]. Beneficial microbes affect specific traits important to plant health and productivity [3]. Consequently, the expression of many important host phenotypic traits relies on host factors and also on the presence and abundance of their associated microbes [4, 5]. Therefore, one of the main goals in the field of plant microbiome research is to improve plant fitness by artificially selecting and manipulating plant-associated microbes that result in the desired host fitness, often referred to as ‘microbiome engineering’.

Several different mechanisms are proposed to be involved in improving microbe-induced plant traits, such as improving plant nutrient uptake [6, 7]), suppression of phytopathogenic microbes [8], modulation of phytohormone production [9] and changes in the plant immune system to mitigate the effects of stress [10]. Through one or a combination of these mechanisms, some important groups of bacteria (e.g., *Bacillus* sp. [11, 12] and *Pseudomonas* sp. [13]) and fungal genera (e.g., *Penicillium* sp. [14, 15] and *Trichoderma* sp. [16, 17]) have been shown to alter quantitative plant traits. Inoculation of plants with single or multiple strains of beneficial microbes has been widely used to enhance plant growth traits, especially for crop plants. Depending on the extent to which the inoculants are efficient, the use of microbial inoculants can contribute towards agricultural productivity with less adverse environmental impacts through the reduction or even elimination of chemical and synthetic fertilizers [18, 19]. However, manipulation of plant microbiomes often does not result in the enhancement of plant traits, and the widespread application of this approach is limited by various biological and experimental factors that affect its efficiency [20]. Regarding biological factors, previous research has shown the advantage of multiple strains over single strain inoculation [21–23]. However, we still have a limited understanding of whether the efficiency of inoculation differs between a wide range of plant models and phenotypic traits, as well as applied microbial phyla. Next to biological factors, the experimental design of inoculation studies may affect the outcome. For instance, the decision on the total time between inoculation and the quantification of plant traits may have an impact on the results and conclusions of a study. Initially, positive effects on plant growth and phenotype may turn negative if these beneficial effects come at some costs [3], leading to the question of the ideal time to detect phenotypic changes in plants. This becomes even more complex when we consider the effect of other important experimental factors on the resulting data, such as inoculation methods, inoculum density, duration of inoculation, and experimental conditions.

Several reviews and meta-analysis studies have been carried out to evaluate the effects of biological (single or multiple strain inoculation [21–23], microbial group [24]) and experimental factors (inoculation method [24], experimental type and conditions [21, 24–26]) on the effectiveness of microbial inoculation on plant traits. Although these studies have contributed significantly to our understanding of the effect of microbial inoculation on plant traits, several important knowledge gaps still need further consideration. To our knowledge, there is no meta-analytic study that considered the influence of a wide range of biological and experimental factors on microbial inoculation results for crop and non-crop plants. This constrains our ability to develop an effective microbial inoculation approach to improve not only traits related to plant biomass (e.g., shoot and root biomass) but also other important traits related to flower and fruit. In terms of experimental factors, there is a lack of information on how the density of the inoculum, the duration of inoculation, and the experimental periods can change the microbial effects on different quantitative phenotypic traits.

To fill these gaps, we performed a meta-analysis to determine how biological and experimental factors shape the outcome of microbial inoculation experiments when the objective is to improve plant phenotypic traits. We reviewed 61 articles covering 27 different plant genera, eight quantitative traits of the plant, and 46 different bacterial and fungal genera, covering a wide range of biological and experimental factors. In addition to discussing how biological and experimental factors influence the effect size of microbial inoculation, we proposed a comprehensive checklist to guide the design, implementation and reporting of microbial inoculation studies. The checklist may standardize microbial inoculation experiments to provide more clear and consistent results leading to more valuable syntheses and more efficient methods to engineer the plant microbiome.

## Material and methods

### Literature search and screening papers

We gathered research papers using the ISI Web of Science and Google Scholar as primary sources. We began our search in May 2021 and ended in February 2022. We used different combinations of keywords such as: (‘microbe’ OR ‘microorganism’ OR ‘bacteria’ OR ‘bacterial’ OR ‘fungi’ OR ‘fungal’) and (‘plant traits’ OR ‘plant growth’ OR ‘plant biomass’). During the first selection round, we checked the titles and abstracts of the studies, excluding studies that did not use microbes to inoculate plants and/or did not measure quantitative traits of plants. The second round of selection involved screening the full texts, with a particular emphasis on material and methods sections of individual papers, to determine whether the studies met the following selection criteria:

I. In each study, the design of the experiment should contain non-inoculated (control) and inoculated (treatment) plants in exactly the same experimental condition.
II. The experiment should test the effect of microbial inoculation on the quantitative traits of the plant, and these data should be reported for both the control and treatment groups. Therefore, to be included in our dataset, a study had to report at least one quantitative trait.
III. Detailed information on biological and experimental factors had to be provided. These factors are listed in the next section on data collection.
IV. If the experiment is performed under certain stress factors (e.g., drought, salinity, heavy metal), the experimental design should contain non-inoculated control and inoculated plants grown under stress and non-stress conditions.
V. Data should be presented so that mean, standard deviation, standard error, and number of replicates could be extracted from the text, tables, and figures.
VI. Research papers had to be published in English as full articles in indexed journals.

The reference lists of identified studies were manually searched using search terms to find additional articles not identified. Finally, 61 studies met the selection criteria and were used in the analysis, which are listed in Supplementary Table S1.

### Data collection

For each study, we extracted the mean, standard deviation, or standard errors, and the number of replicates corresponding to specific plant traits in the control and treatment groups to calculate effect sizes. In cases where standard error (SE) was reported, this was converted to standard deviation (SD) by the equation: 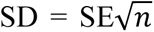, where *n* is the number of replicates. Our data extraction resulted in the following variables for biological factors: (1) overall effect size of microbial inoculation, (2) plant trait categories, (3) plant models (crops and non-crop plants), (4) single or multiple species inoculation, (5) microbial phyla and (6) microbial sources. To quantify the effect of microbial inoculation on plant traits, we extracted the following eight plant trait categories in our data set: (a) overall plant growth (plant fresh and dry weights and plant height), (b) shoot (fresh and dry weight, length, branch number), (c) root (fresh and dry weight, length), (d) leaf (leaf area, leaf number, fresh and dry weight, leaf chlorophyll content), (e) flower (flowering time), (f) fruit (fruits and pod number and weights), (g) nodulation (nodules fresh and dry weights and nodules number), and (h) seeds (seed germination and seed numbers or yield). Microbial sources indicate the setting in which microbes were extracted, including soil, rhizosphere, seed and leaf, root, plant endophytes (not specified), and commercial microbes.

As for experimental factors, we collected information about (1) seed sterilization, (2) inoculation methods, (3) inoculum density (low, medium, and high density, see below), (4) with or without stress application, (5) experimental conditions, and (6) experimental periods. The seed sterilization category indicates whether or not plant seeds were sterilized before inoculation with microbes. Inoculation methods contain information about which part of the plant has been inoculated, including seed, seedlings, root zone (root or rhizosphere treatments), and culture or potting growth medium. Seed inoculation refers to a process where liquid inoculants are prepared and applied to the seed. The density of the inoculum determines the range of microbial cells in the solution used to inoculate the plants. Based on the value obtained from the studies, we classified the inoculum density into four groups that ranged from low (1 ×10^6^ - 1 ×10^7^ CFU or spores/ml or gr/soil), medium (1 ×10^8^), high (1 ×10^9^ - 1 ×10^10^) or ‘not specified’ for the studies that did not report these values. Experimental conditions indicate whether the experiment was conducted under sterilized (meaning that substrates that plants grow in, such as soil, sand, or agar, were sterilized before the addition of microbes) or unsterilized growth substrates or under field conditions. We also noted the duration from plant inoculation to trait measurement as experimental periods.

### Statistical analysis

All statistical analyses were performed in R environment 4.2.2 (R Development Core Team 2021). We performed meta-analysis using the “metafor” package [27]. This R package has been commonly used in previous meta-analysis studies [21, 24, 28]. For each observation, we calculated the effect sizes (i.e., the magnitude of the microbial inoculation effect on plant traits) based on the log response ratio (lnRR) using the ‘ROM’ option (the log-transformed ratio of means; [29]) with the escalc() function. The formula is *lnRR = ln(X_t_/X_c_)*, where *X_t_* and *X_c_* are the mean values in inoculated and non-inoculated plants, respectively. The variance of lnRR was calculated as follows: *V _lnRR_ = (SD X_t_)^2^/n_xt_ (X_t_)^2^ + (SD X_c_)^2^/n_xc_ (X_c_)^2^* where *SD X_t_* and *SD X_c_* are the corresponding standard deviations and *n_xt_* and *n_xc_* are the sample size in inoculated and non-inoculated plants, respectively. We selected lnRR as our effect size metric because the log transformation of parameters reported in different units across studies better supports normal distribution for statistical analysis [30]. This is particularly important when different units have been used to measure plant traits within and between studies, which is the case in our study.

We performed a linear mixed effect model (a multilevel meta-analysis) using ‘metafor:: rma.mv’ with paper IDs as random factors, lnRR as the response variable, and the moderators (that is individual biological and experimental variables that may shape the magnitude of the effect size) as fixed effects. Because most studies contained multiple observations, we included the ID of the paper as a random effect in our analysis [31]. We used restricted maximum likelihood (REML) tests to assess the homogeneity of variances within the study and between the studies [27, 32]. We checked for publication bias using the Rosenberg failsafe number [33] and tested the asymmetry (funnel plot) of effect sizes using an ‘rma.mv’ model with the function ‘mod=vi’ for the entire dataset. Then, we first estimated an overall effect size (or overall mean) that included only an intercept and the paper IDs as a random effect without considering the effect of moderators (variables). To address which biological and experimental factors (as moderators) individually influence the effect size, we separately added each of the eleven moderators as fixed effects to our base model described above (11 models in total). The estimated effect sizes of the meta-analysis, together with their 95% confidence intervals (CI) and prediction intervals (PIs - which show heterogeneity between effect sizes), were shown based on orchard plots using the ‘orchard’ package [34].

To account for the level of precision between studies, the individual effect size was weighted according to the variance of lnRR and the number of replicates (n), thus giving more weight to well-replicated studies and those with low SD values. On top of that, since our meta-analyses included multiple moderators in such a way that each has several sublevels, the effect size is further weighted based on “marginalised” means (using weights = “prop” function) in orchard plots. This is very important in case of unequal sample size in categorical variables (which is the case in the majority of biological and experimental variables in our dataset). Therefore, for each subgroup within each moderator, this function weights the effect size based on its proportional representation in the data. For this reason, in some cases (see Fig. 1B for flower in plant trait categories and Fig. 2F for more than 3 months in the experimental period), the weighted mean effect size seems out of the range in orchard plots but is still correct (see here [34] for more explanation). If the effect was statistically significant (p-value < 0.05), we performed pairwise comparisons of subgroups within each moderator level using Tukey’s HSD post hoc test based on the function ‘ghlt’ using the package ‘multcomp’ [35]. The mean effect size (for subgroups within each level of a moderator) was considered significant when the confidence interval did not overlap with zero. The estimates effect size and 95% CI used in post hoc comparisons and in figures were obtained from models that included only interactions compared to a zero intercept, without the main effects [36].

**Figure 1.**
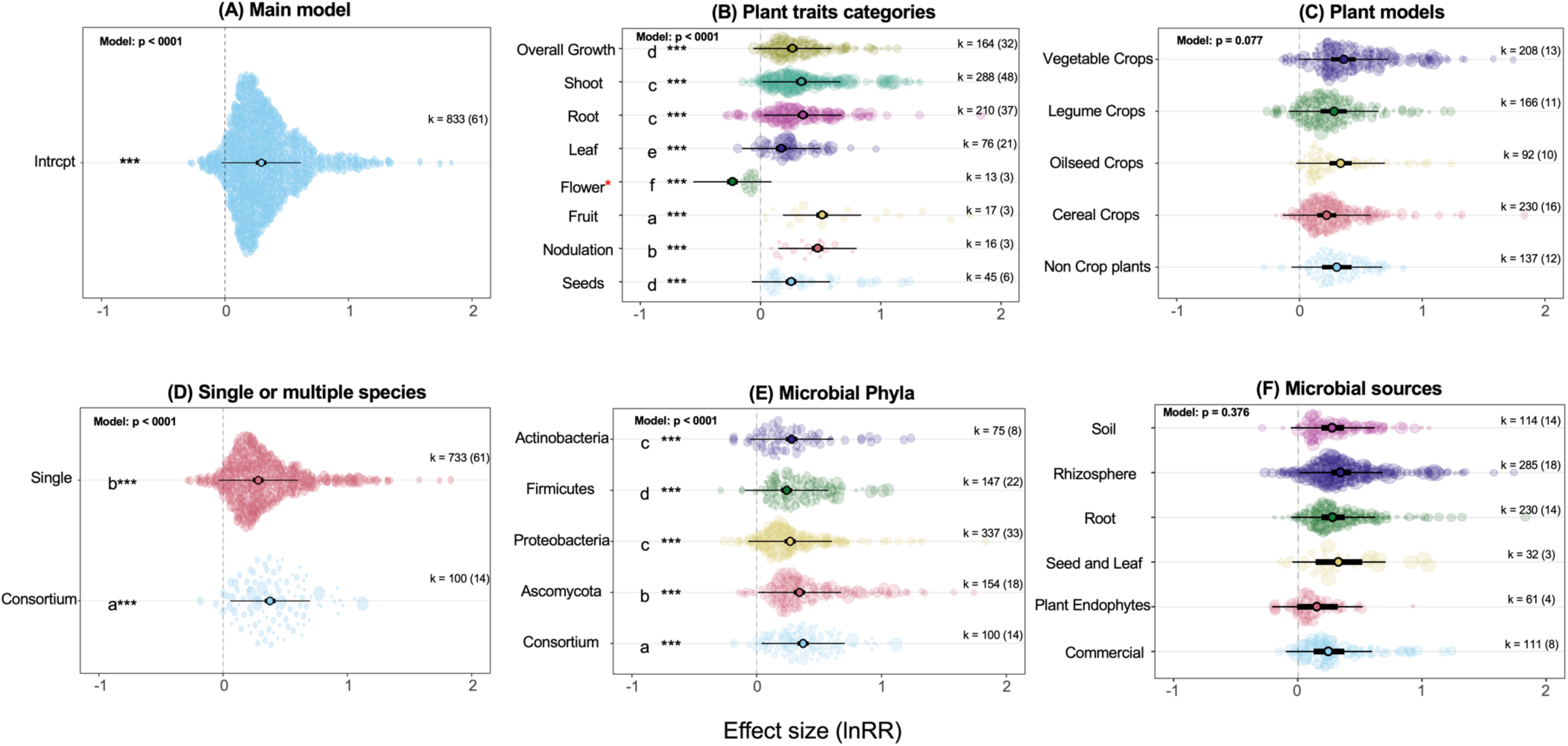
The estimated overall effect size (lnRR) of microbial inoculation on the quantitative traits of the plant (**A)**. The individual effect of biological factors on the strength of the effect size (**B-F**). Solid dots with thick black bars represent the effect size and 95% confidence intervals (CI). The thin bars are representative of the prediction intervals (PI), indicating the heterogeneity among the effect sizes or expected values that future studies may find. The transparent, colorful circles show individual data points whose size is adjusted according to the precision of the study. The size of each colorful circle is proportional to its relative weighting in the overall model and the number of replicates (n), thus giving more weight to well-replicated studies. The k-value represents the number of observations (data points) in the model, while the value in parentheses shows the number of studies. If the CIs do not cross the vertical dotted lines (lnRR = 0), the effect size is significant at p < 0.05. If the effect size was statistically significant for each biological factor (at p < 0.05), we performed pairwise comparisons of subgroups within each biological factor using Tukey’s HSD post hoc test. Only for those significant biological factors are shown, asterisks that are denoted to sublevels that are significant within each biological factor (if the CIs do not cross the vertical dotted lines). Letters represent significant pairwise differences between sub-levels based on post hoc Tukey’s test. * For the flower, only the number of days required for flowering was included in this sub-category. Therefore, the significant negative effect size means that the plants inoculated with microbes flowered earlier than the non-inoculated control plants. See the main text for more explanation.

**Figure 2.**
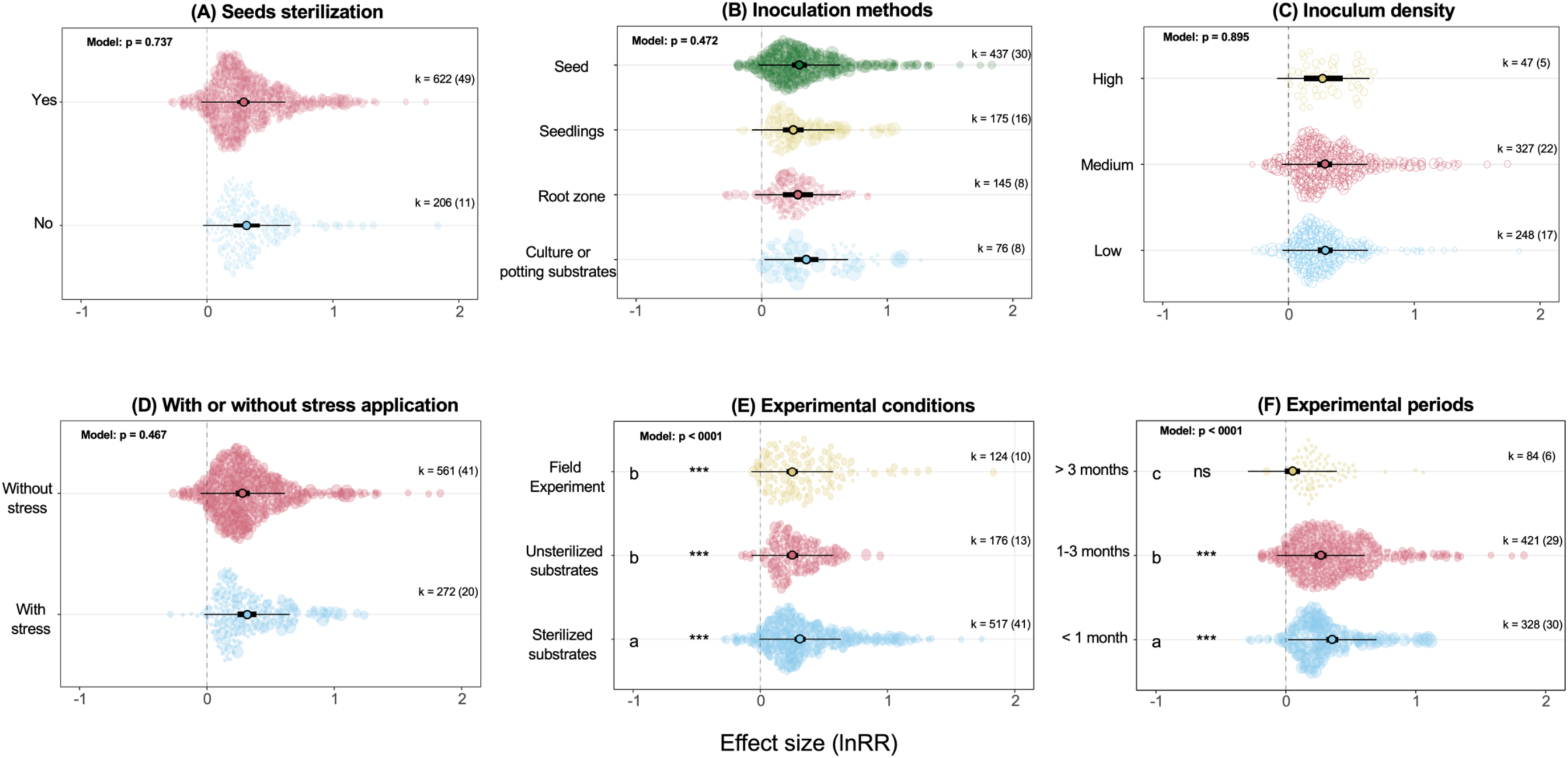
The individual effect of experimental factors on the strength of the effect size. Asterisks are denoted to sub-levels that are significant within each biological factor, while letters represent significant pairwise differences between sub-levels based on post hoc Tukey’s test. See Figure 1 for a general explanation of the Orchid plot.

## Results

We reviewed 61 articles, which resulted in 833 individual observations (effect sizes), covering 27 different plant genera, eight trait categories, and 46 different bacterial and fungal genera (Supplementary Table S1). The rank correlation test indicated a moderate asymmetry in the funnel plot, suggesting some evidence of publication bias (Kendall’s τ = 0.0487, p = 0.035). However, the estimation of the Rosenthal failsafe number, which estimates how many additional effect sizes with no effect (non-significant studies) would be necessary to alter the observed effect size in our study, suggests that our findings are unlikely to be due to chance (Failsafe N = 10,563, average effect size = 0.23, observed significance level < 0.0001).

### Which biological factors influence the effect size of microbial inoculation?

During the course of data extraction from selected articles, we identified and extracted different biological factors that could explain the strength of the effect size. These biological factors and their sublevels are shown in Figure 1. Overall, we observed that the inoculation of plants with microbes had a positive effect size than non-inoculated control plants (lnRR = 0.29, 95 % CI = 0.24–0.33, p < 0.0001; Fig. 1A). Within our meta-analysis, the traits related to the shoot were the most studied traits (34.4% of total observation) compared to the traits of fruits (2%), nodulation (2%), and flowers (2.1%) traits (Fig. 1B). Depending on the plant trait categories, the magnitude of the effect sizes varied significantly (p < 0.0001; Fig. 1B), with traits related to fruit (lnRR = 0.50, 95 % CI = 0.46–0.55), nodulation (lnRR = 0.47, 95 % CI = 0.42–0.52), root (lnRR = 0.35, 95 % CI = 0.30–0.39), and shoot (lnRR = 0.33, 95 % CI = 0.29–0.38), being most positively influenced by microbial inoculation. As for the flower, we only included the number of days required for flowering in this sub-category. Therefore, the significant negative effect size (lnRR = −0.23, 95 % CI = −0.28 – −0.19) means that the plants inoculated with microbes flowered earlier than the non-inoculated control plants, thus, indicating that microbes impose a positive effect on flowering time. The plant model did not significantly influence the effect size (p = 0.077; Fig. 1C). Inoculating plants with the microbial consortium (meaning that more than one microbial species were included in inoculation) had a greater positive impact on effect size (lnRR = 0.37, 95 % CI = 0.33–0.42) than inoculation with a single microbial species (lnRR = 0.28, 95 % CI = 0.28–0.32; Fig. 1D). Furthermore, the microbial phyla significantly influenced the strength of the effect size (p < 0.0001; Fig. 1E). The most dominant phyla studied were *Proteobacteria* than other microbial phyla (Fig. 1E). Although all microbial groups showed significant positive effects on overall effect size (p < 0.0001; Fig. 1E), *Ascomycota* (as fungi phyla) appeared to insert a greater effect (lnRR = 0.34, 95 % CI = 0.29–0.38) in compared to bacterial phyla (Fig. 1E). Similar to the plant model, the microbial sources (p = 0.376; Fig. 1F) did not significantly influence the effect size.

### Which experimental factors influence the effect size of microbial inoculation?

To further explore the potential effect of experimental factors on the effect size of microbial inoculation on plant traits, we focused on six factors, which are shown in Figure 2. Seed sterilization, inoculation methods, inoculum density, and whether plants were under stress conditions or not did not have a significant effect on the estimated effect size (Fig. 2A-D). As for inoculum density, the majority of the studies (39%) did not report these values and were categorized as “Not Specified” (which is not shown in the figure and not included in the main test). The experimental conditions significantly affected the effectiveness of microbial inoculation in plant traits (p < 0.0001; Fig. 2E). The largest estimated lnRR (lnRR = 0.31, 95 % CI = 0.27–0.35) was observed when the experiments were carried out on sterilized substrates (meaning that substrate in which plants grow was sterilized before the addition of microbes). On the other hand, the effect size became smaller but still positive and significant when plants were grown in unsterilized substrate and field experiments (lnRR = 0.253 and 0.250, respectively). We further investigated the impact of experimental periods (from the start of the experiment until plant traits were measured) on effect sizes. A significant portion of the data sets analyzed conducted the experiment within a month (49%), and only 9.8% of the studies were performed more than three months, and the remaining studies were within these periods (Fig. 2F). Our results showed that the experimental periods had a significant effect on the effect size: the strongest positive effect was observed for those experiment performed within a month (lnRR = 0.35, 95 % CI = 0.31–0.40), and the lowest non-significant effect for those more than three months (lnRR = 0.05, 95 % CI = −0.008 –0.109; Fig. 2F).

## Discussion

Our meta-analysis showed that inoculating plants with multiple bacterial and/or fungal strains significantly influences quantitative plant traits more than a single isolate inoculation. We revealed that the plant trait under consideration and the microbial taxonomic identity significantly shape the outcomes of microbial inoculation experiments. In contrast, the type of host plant (e.g., crop or non-crop plant), as well as the source of microbes (e.g., soil or phyllosphere), do not explain variation in inoculation effect size. Traits related to fruit, nodulation, root, and shoot showed stronger responses to microbial inoculation than leaf, flower, and seed traits. Next to these biological factors, the experimental design of inoculation studies also impacts their outcome in such a way that experiments under sterilized conditions and those that terminate after relatively short periods resulted in larger positive effects on plant traits than those of unsterilized and long-term experiments. In general, these findings provide valuable insight into the effects of biological and experimental factors on the outcomes of microbial inoculation on plant traits. Understanding which plant traits are most responsive, the differential effects of microbial taxonomic groups, and the potential benefits of microbial consortia can help to design optimal microbial inoculation treatments that lead to robust results under variable environments, including field conditions.

### Biological factors influencing the effect size

Our results revealed that the significant variation in effect sizes depends on the identity of plant traits, highlighting the importance of a trait-specific understanding of plant-microbe interactions. For instance, traits related to root and nodulation were among those that were strongly influenced by microbial inoculation. Since inoculants are often applied to the root environment, roots are the primary plant part that interacts with inoculated microbes in these cases, which may lead to stronger effects of microbial inoculation on root traits observed in our study. For the nitrogen-fixing nodulation trait, legume plants actively select and attract nitrogen-fixing rhizobial strains by supporting the growth and attracting them towards themselves by producing and releasing specific root exudate compounds (e.g., flavonoids) [37]. Collectively, these observations highlight the specificity of plant-microbe interactions.

Most of the studies in our analysis applied single-strain inoculation rather than multiple-strains, where *Penicillium* sp. and *Trichoderma* sp. were the most dominant fungi genera and *Bacillus* sp. and *Pseudomonas* sp. being the most widely used bacterial genera. The fungal phylum *Ascomycota* exhibited a larger effect than the bacterial phyla when only single-strain inoculation was considered. One possible explanation for the greater effect of *Ascomycota* than bacterial phyla can be due to their multi-directional beneficial effects on plant traits. Previous research has shown the direct effects of *Penicillium* spp. on enhancing plant growth phytohormones (e.g., indole acetic acid (IAA) and siderophore production [14]; Gibberellic acids [15]) and solubilization of phosphates [14]. *Trichoderma* spp. has been widely reported to significantly inhibit pathogenic plant microbe growth (indirect effects) and improve plant growth traits directly via the production of secondary metabolite compounds (see review on this topic here [16, 17]). In addition to these beneficial effects, many members of fungi in the *Ascomycota* phylum are known to form extensive hyphal networks in the soil, which can facilitate plant nutrient and water uptake, thus directly affecting plant growth and thus traits. Furthermore, some groups of bacteria take advantage of hyphal networks that facilitate their dispersal network [38], eventually helping them to occupy the plant environment. The combination of these beneficial effects may explain the observed differences in effect size between *Ascomycota* and bacterial phyla.

We showed that inoculation of plants with microbial consortia is more effective than single-strain in improving plant traits, supporting previous metadata studies [21, 22]. A possible explanation for a larger effect of consortium than single strain inoculation can be due to the variable and synergistic plant growth-promoting activities that co-inoculation of more than one strain can offer to plants in comparison with single-strain inoculation. For example, co-inoculation of *Bradyrhizobium* sp. and *Leclercia adecarboxylata* improved a wide range of plant traits, such as longer roots, higher nodulation rates, higher shoot nutrient contents (e.g., N and P) and yield than in single inoculation or not inoculated soybean plants under field conditions [9]. In another example, Jain et al [39] showed that plants (*Pisum sativum*) treated with a combination of three microbes as a consortium (for bacteria: *Pseudomonas aeruginosa* and *Bacillus subtilis*; for fungi: *Trichoderma harzianum*) enhanced plant growth characteristics (e.g., plant length and biomass) in the presence and absence of *Sclerotinia sclerotiorum* as a plant pathogen. Beneficial bacterial strains may take advantage of fungi mycelia, which would support their dispersal to have a better chance of occupying the plant environment [38]. Therefore, such collaborative interactions between multifunctional microbial strains may increase the chance that the inoculated consortium better establishes and colonizes the plant environment [20].

### Experimental factors influencing the effect size

Our data did not show significant differences in effect sizes among seed, seedling, root, and soil inoculation, which is consistent with previous studies showing that inoculation methods do not change the output of the microbial effect on plant performance [21]. A larger effect size was observed when experiments were performed under sterilized conditions than when plants were inoculated under unsterilized or field conditions. These results confirmed previous meta-analysis studies [21, 25], in which they reported a significantly larger effect size in the pot or greenhouse experiment compared to field conditions. One possible explanation for these findings is that sterilizing the substrates that are used for plant growth may provide better conditions for inoculated microbes to establish themselves and colonize the plant environment. This can occur because of the absence (or reduced diversity) of other microbes; thus, inoculated microbes do not have competitors under such experimental conditions, which can magnify the effects of inoculation treatment [20]. Therefore, while growing the plant in a sterilized pot or agar media provides valuable information on the effect of microbes on plant traits, they may not be fully representative of the growth conditions that plants face in natural settings [40]. It is also important to note that the successful establishment of microbes in experimental settings was not tracked in most of the studies included in our analysis. Thus, we do not know how persistent the inoculated microbes are. Future studies should monitor the persistence of inoculated microbes and their interactions with native microbes over the course of the experiment.

In terms of the duration of the experiment, we showed that short-term experiments (those studies that were carried out in less than a month) resulted in the strongest positive effect compared to mid-term experiments (1-3 months) and long-term experiments (more than 3 months). Short-term experiments capture only the more pronounced effects of microbial inoculation (the initial phase), which can be partly due to immediate changes in the availability of nutrients in the plant environment that may improve some specific plant traits. The decrease in positive effects during mid-term experiments may indicate that the initial strong response diminishes over time, which can be indicative of plant adjustments to the microbial community, leading to a stabilization of effects. When plants adjust the microbial populations in the long run, it may lead to non-significant effect sizes of microbial inoculation. An example of this is reported for tomato plants where inoculated bacterial strains appeared to significantly enhance traits related to the seedling’s aboveground biomass under short-term greenhouse experiments (30 days) [41], while using the same strains resulted in a non-significant effect on fruit traits (e.g., number of fruits per plant and fruit weight) under long-term semi-field conditions [42]. Therefore, the observed enhanced effect of microbial inoculation in the short term may not necessarily correspond to a better performance at the harvest stage (e.g., seeds or fruit numbers).

Short-term studies in which host plants have not reached their final growth stages neglect the possible (positive or negative) microbial effect in the final stages of plant development, which is of high importance for crop plants as it corresponds directly to agricultural productivity. A field experiment previously showed that the plant’s development stages play an important role in shaping bacteria and fungi communities associated with the rhizosphere, roots, and leaves in wheat [43]. In light of these observations, it becomes clear that although the short-term benefits of microbes on some plant traits are well documented, our understanding of their long-term effects remains a critical knowledge gap. To address this gap and provide more comprehensive insights, future research should prioritize conducting experiments that extend to the later phases of host plant growth and development (Fig. 3). These studies should not only explore the immediate or intermediate impacts of inoculated microbes on plant phenotypes but also assess how these effects evolve over time, as well as interactions with soil or plant-inhabiting microbes. New phenotyping technology that would allow scanning the plant and capturing various morphological and physiological traits in a non-destructive way are promising tools for plant phenotyping [44]. Such tools will allow us to perform a series of time-point measurements over the course of microbial inoculation experiments, thus better tracking plant-microbe interactions.

**Figure 3.**
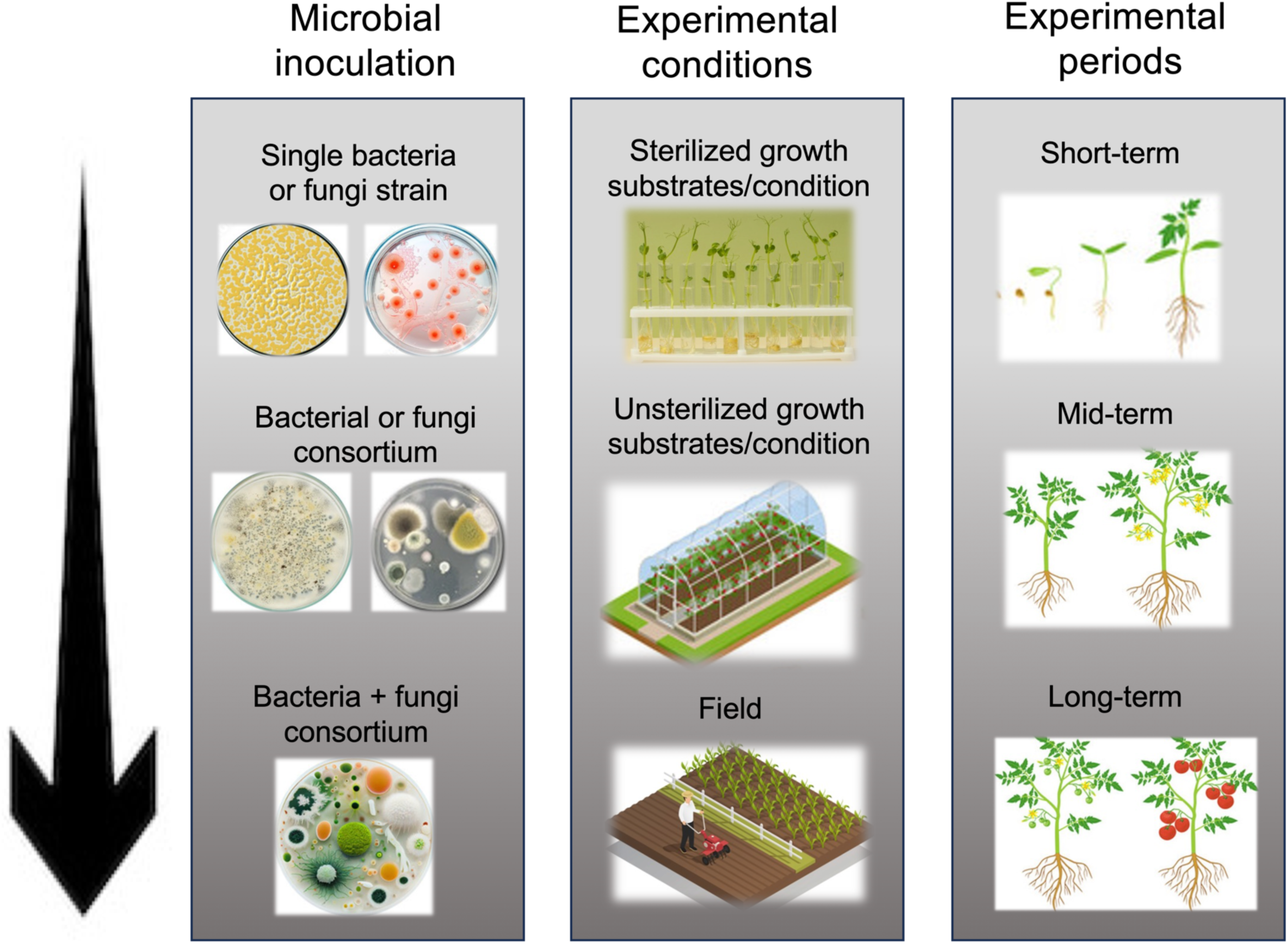
Future research directions in using microbes to improve plant traits. In term of microbial inoculation, we recommend that future studies should consider inculcating plants with a consortium containing both bacteria and fungi strains instead of a single strain. The short- and long-term effects (experimental periods) of such microbial consortiums on plant traits should be tested not only under sterilized and non-sterilized growth conditions but also under robust field experiments (experimental conditions) with naturally occurring microbial communities associated with the plants.

### Checklist for microbial inoculation experiments

In general, we did not identify a standard framework for conducting microbial inoculation experiments in the studies included in our meta-analysis, but experimental designs were mostly study-specific. In addition, important experimental conditions were not consistently reported, which calls for standardized protocols. We found many papers fit in the scope of the study but were ultimately not included in the meta-analysis due to the lack of important information. For example, in many cases, the authors did not specify standard error/ deviation or number of replicates or did not provide sufficient information about biological factors and experimental settings. Standard protocols that include details on the biological and experimental factors that we discussed are essential to replicate and reproduce the patterns observed in each study and also to synthesize individual studies to general knowledge. Therefore, here we proposed a checklist, including some guidelines and recommendations for microbial inoculation experiments, with the aim of providing future studies with a structured framework to design, conduct, and report their experiments (Table 1). This checklist encompasses key elements for microbial inoculation-based studies, including microbial strains, plant models and traits, environmental conditions, and data reporting. By following these guidelines, experiments will become more transparent, making it easier for others to replicate our work and facilitate more robust meta-analyses. Thus, including a wider range of studies in future syntheses will be possible by following this approach. Additionally, this checklist can serve as a valuable resource for journal editors and peer reviewers, allowing them to evaluate the completeness and transparency of submitted manuscripts in the context of microbial inoculation research.

**Table 1.** Checklist and recommendations for microbial inoculation experiments.

|  |  |
| --- | --- |
| Factors | Notes |
| Microbes | **<u>Identification of microbial isolates:</u>**<br>○ Detailed information about microbial strains, including strain/species name and their corresponding sequences.<br>**<u>Source of microbial isolates:</u>**<br>○ Specify the source from which microbial candidates were isolated.<br>○ If microbes were isolated from the plant compartment, report the plant's identity and the plant compartment as well as their development stage and the environmental conditions when sampling was performed to isolate microbes.<br>**<u>Inoculum preparation:</u>**<br>○ Details on preparing and handling microbial inoculum (culturing in liquid or solid media, type of media, etc.).<br>○ Specify the exact density of the microbial cell included in the inoculation. This should be reported for each strain if multiple strains are included in the treatment. Use a standard way for reporting such data (e.g., CFU/ml for bacteria or spores/ml for fungi). |
| Plant model and traits | **<u>Information on plant model:</u>**<br>○ The detailed information about the plant model, including species/cultivar name.<br>○ Specify the source where plant seeds were provided.<br>**<u>Plant trait measurements:</u>**<br>○ Detailed information on measured plant traits. The best would be to include more than one trait as a response variable.<br>○ Report the methods used for trait measurements.<br>○ Indicate the units for each plant trait. |
| Experimental settings | **<u>Inoculation methods and duration:</u>**<br>○ Whether or not plant seeds were surface sterilized before starting the experiment.<br>○ Specify the inoculation methods. This can include information on which exact plant parts (seeds, seedlings, root zone (root or rhizosphere treatments)), or culture or potting growth media microbes were introduced.<br>○ Specify the duration of microbial inoculation in cases where seeds or part of seedlings (e.g., roots) have been immersed or soaked in microbial solution before plants were transferred to the pot.<br>**<u>Experimental setup and conditions:</u>**<br>○ Specify whether the experiment is performed in the pot (with sterilized or unsterilized potting media), agar, or under greenhouse or field conditions.<br>○ Details on plant growth conditions, such as light intensity, temperature, and humidity.<br>○ In case plants are grown in soil, specify soil characteristics (organic matter content, pH, and nutrient content or product name in case of commercially available soils).<br>○ In case plants are grown in agar media, specific types of agar media (e.g., LB, R2A, etc.).<br>○ Experimental period (from inoculation to the trait measurement).<br>○ Specify the geographical location where the experiments were conducted. |
| Data reporting | ○ Use standardized units and measurement scales for consistency.<br>○ For each variable, report the mean, standard error/deviation, and the number of replicates (preferably in a table format). Share plant trait data (and other relevant information) in a repository and follow FAIR principle [45].<br>○ Share detailed information on the statistical methods used. |

### Concluding remarks and future research directions

Plant-associated microbes are an important player in affecting plant traits, and thus, microbiome engineering is a promising approach to increase plant performance. We have presented evidence of how plant traits are altered by microbial inoculation, highlighting the importance of trait-specific interactions, the differential effects of microbial taxonomic identity, and the potential benefits of microbial consortia, which can all contribute to altering plant phenotypes. Our findings provide new insight into the importance of biological and experimental factors that shape the effects of microbial inoculation on plant phenotypes. These insights may contribute to the effective identification of microbial candidate strains that persist in natural settings and improve plant growth and function, offering exciting prospects for sustainable agriculture in a rapidly changing climate. Future research should optimize microbial inoculation effect on plant traits by varying biological and experimental parameters summarized in Figure 3. From inoculating plants with single bacteria or fungi strains, we need to move towards using consortia that include both bacteria and fungi. The short- and long-term effects of such microbial consortia on plant traits should be tested not only under sterilized growth conditions but, more importantly, also in (semi-)field experiments. The results of such studies may contribute to the effective use of microbial inoculants and maximize the microbial effect to improve plant growth and functioning under variable environments.

## Acknowledgments

We would like to thank the members of the Evolutionary Ecology of Plants research group at Philipps-University Marburg for the fruitful discussion, which greatly helped to build the ideas behind this study.

